# ER-phagy Receptor’s Intrinsically Disordered Modules Drive ER Fragmentation and ER-phagy

**DOI:** 10.1101/2024.06.18.599470

**Authors:** Mikhail Rudinskiy, Carmela Galli, Andrea Raimondi, Maurizio Molinari

## Abstract

Membrane remodeling leading to fragmentation is crucial for autophagy programs that control capture by phagophores or endolysosomes of portions of organelles to be removed from cells. It is driven by membrane-bound autophagy receptors that display cytoplasmic intrinsically disordered modules (IDRs) engaging Atg8/LC3/GABARAP (LC3). Studies on endoplasmic reticulum (ER)-phagy receptors of the FAM134 family revealed the importance of sequential FAM134 proteins phosphorylation, ubiquitylation and clustering for execution of the ER-phagy programs. In this model, ER fragmentation is promoted/facilitated by the membrane-remodeling function of FAM134 reticulon homology domains (RHDs). However, RHDs are not conserved in ER-phagy receptors. The question that we tackle in this work is if activation of ER-phagy receptors anchored at the ER membrane with conventional membrane spanning domains, i.e., most of the ER-phagy receptors known to date, eventually trigger ER remodeling and fragmentation, and how. Here, we show that the membrane-tethering modules of ER-phagy receptors (RHDs for FAM134B, single/multi spanning transmembrane domains for TEX264 and SEC62) determine the sub-compartmental distribution of the receptors but are dispensable for ER fragmentation, regardless of their propensity to remodel the ER membrane. Rather, ER fragmentation is promoted by the ER-phagy receptors intrinsically disordered region (IDR) modules that are a conserved feature of all ER-phagy receptors exposed at the cytoplasmic face of the ER membrane. Since cytoplasmic IDRs with net negative charge are conserved in autophagy receptors at the limiting membrane of other organelles, we anticipate that conserved mechanisms of organelle fragmentaVon driven by cytoplasmic exposed IDRs could operate in eukaryoVc cells.

## Introduction

ER-phagy regulates size and activity of the ER, the organelle that produces proteins, lipids and sugars in nucleated cells. ER-phagy dysfunction is linked to muscle, heart, liver diseases, neurodegeneration, ER storage disorders, and defective biogenesis of organs and tissues. Activation of ER-phagy due to elevated expression of ER-phagy receptors is observed in quickly progressing cancers (Bergmann *et al*, 2017; Chipurupalli *et al*, 2022; Hagerstrand *et al*, 2013; Zimmermann *et al*, 2022). ER-phagy receptors control the fragmentation and lysosomal clearance of ER portions that are damaged, aged, superfluous, or contain toxic macromolecules. The ER-phagy receptors are all composed of a membrane-tethering module and of a cytoplasmic intrinsically disordered region (IDR) containing a LC3 interacting motif (LIR) (Adriaenssens *et al*, 2022; Chino & Mizushima, 2023; Conway *et al*, 2020; Ferro-Novick *et al*, 2021; Hubner & Dikic, 2020; Johansen & Lamark, 2020; Lamark & Johansen, 2021; Mizushima, 2022; Popelka & Uversky, 2022; Reggiori & Molinari, 2022). Seminal studies performed to understand the mode-of-action of mammalian ER-phagy receptors of the FAM134 family (FAM134A, FAM134B, FAM134C) established that their phosphorylation, ubiquitylation, and homo-/hetero-oligomerization control ER fragmentation and lysosomal clearance (Berkane *et al*, 2023; Di Lorenzo *et al*, 2022; Foronda *et al*, 2023; Gonzalez *et al*, 2023; Iavarone *et al*, 2022; Jiang *et al*, 2020; Khaminets *et al*, 2015; Reggio *et al*, 2021). FAM134 proteins are peculiar ER-phagy receptors because they possess a RHD (light blue in **Fig. 1A**), a membrane-tethering module found in ER shaping proteins of the reticulon and REEP families that generates high curvature and constrictions of the ER membrane (De Craene *et al*, 2006; Hu *et al*, 2008; Hu *et al*, 2009; Kiseleva *et al*, 2007; Shibata *et al*, 2010; Shibata *et al*, 2008; Terasaki *et al*, 2013; Tolley *et al*, 2008; Voeltz *et al*, 2006; Wang *et al*, 2013; Wu & Voeltz, 2021). The capacity of the FAM134B RHD to remodel the ER membrane has been confirmed by molecular dynamics simulations and *in vitro* liposome remodeling (Bhaskara *et al*, 2019) and has been proposed to promote/facilitate the ER fragmentation required for lysosomal clearance of selected ER portions (Bhaskara *et al*., 2019; Foronda *et al*., 2023; Gonzalez *et al*., 2023; Gubas & Dikic, 2022; Hoyer & Harper, 2023; Jiang *et al*., 2020; Khaminets *et al*., 2015; Siggel *et al*, 2021). However, membrane remodeling is not a conserved trait of membrane-tethering modules of ER-phagy receptors. In fact, the majority of Fungi, Plant and Metazoan ER-phagy receptors are anchored via conventional multi- or single-spanning transmembrane domains that are not expected to promote membrane remodeling *per se* (TMD, blue in **Fig. 1A** for the mammalian ER-phagy receptors SEC62 and TEX264). Moreover, ER-phagy receptors have been identified that lack membrane anchor and are recruited at the ER membrane only upon activation of the ER-phagy programs (Reggiori & Molinari, 2022). Yet, activation of all ER-phagy receptors (independent of their mode-of-anchoring to the membrane or of their soluble status) triggers the ER fragmentation that obligatorily precedes lysosomal delivery. Since the common trait of ER-phagy receptors is the presence of long cytoplasmic IDR modules, we postulate that these modules play an active role facilitating ER fission.

**Figure 1.**
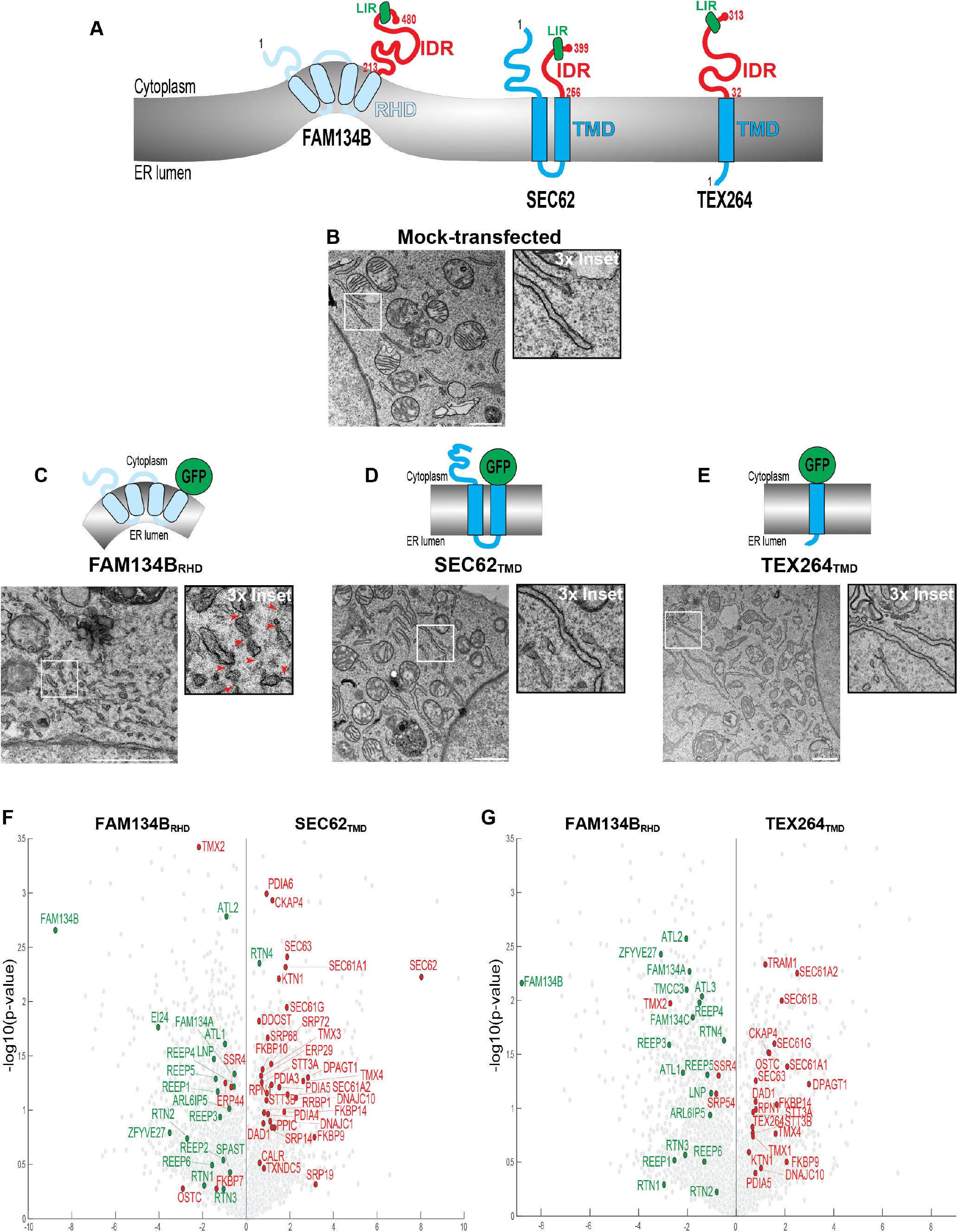
Membrane remodeling and sub-compartmental localization functions of membrane-tethering modules. **(A)** ER-phagy receptors FAM134B, SEC62, and TEX264. TMD: transmembrane domain; RHD: reticulon homology domain; LIR: LC3-interacting region; IDR: intrinsically disordered region. **(B)** RT-TEM showing ER ultrastructure in mock-transfected MEF. Scale bar: 1 μm (see **Movie S1** for Room Temperature-Electron Tomography, RT-ET). **(C)** Same as (B) showing constricted ER tubuli in MEF expressing FAM134B_RHD_ and **Movie S2. (D)** Same as (B) for SEC62_TMD_-transfected cell and **Movie S3. (E)** Same as (B) for TEX264_TMD_ and **Movie S4. (F)** Vulcano plot, endogenous ER proteins among FAM134B_RHD_ or SEC62_TMD_ interactors in HEK293 cells, N=2 biological replicates. **(G)** Same as (F) for FAM134B_RHD_ and TEX264_TMD_.

Here, we assess the capacity of ectopically expressed ER-phagy receptors and of their membrane-tethering and cytoplasmic IDR modules to recapitulate activation of ER-phagy programs that rely on fragmentation of the ER followed by delivery of the ER fragments to endolysosomal compartments for clearance. We show that ectopic expression of FAM134B, SEC62 and TEX264 triggers ER-phagy, even though only the membrane-tethering module of FAM134B shows the capacity to remodel the ER membrane. The membrane-tethering modules of FAM134B, SEC62 and TEX264 determine the receptor’s distribution within the ER but are dispensable for ER fragmentation. Crucially, the expression of the ER-phagy receptors’ IDRs tethered at the ER membrane with a short sequence of hydrophobic amino acids is sufficient to trigger ER fragmentation, and to deliver ER portions to the degradative compartments if the IDRs engage LC3. Cytoplasmic IDRs engaging LC3 proteins are functionally conserved amongst membrane-bound autophagy receptors that control fragmentation/fission for lysosomal clearance of other organelles such as mitochondria or the Golgi compartments. We therefore anticipate the existence of conserved mechanism of organelle fragmentation relying on the activajon of IDRs at the organellar limiting membrane.

## Results

### ER membrane remodeling by membrane-tethering modules

To elucidate the role in ER-phagy of the membrane-tethering and of the cytoplasmic IDRs modules composing the ER-phagy receptors (blues and red, respectively, **Fig. 1A**), we first monitored in electron microscopy the ER morphology in mouse embryonic fibroblasts (MEF) expressing the RHD of FAM134B, the double-spanning TMD of SEC62, or the single-spanning TMD of TEX264. To this end, MEF were mock-transfected (**Fig. 1B**), or were transfected with plasmids for expression of FAM134B_RHD_ (**Fig. 1C**), SEC62_TMD_ (**Fig. 1D**), or TEX264_TMD_ (**Fig. 1E**), where the cytoplasmic C-terminal IDRs of the three ER-phagy receptors (red in **Fig. 1A**) is replaced with the monomeric superfolder (sf)GFP (Pedelacq *et al*, 2006). Room Temperature-Transmission Electron Microscopy (RT-TEM, **Fig. 1C**) and RT-Electron Tomography (RT-ET, **Movie S2**) show that the expression of FAM134B_RHD_ dramatically changes the morphology of the ER (compare with mock-transfected cells, **Fig. 1B** and **Movie S1**), with the generation of thin constrictions (red arrowheads in **Fig. 1C**, Inset) and highly tubular interconnected ER network. The same modification of the ER ultrastructure has previously been reported upon expression of other reticulon and REEP family members, whose expression generates and constricts ER tubuli *in vitro* and *in cellula (De Craene et al*., *2006; Hu et al*., *2008; Hu et al*., *2009; Kiseleva et al*., *2007; Shibata et al*., *2010; Shibata et al*., *2008; Terasaki et al*., *2013; Tolley et al*., *2008; Voeltz et al*., *2006; Wang et al*., *2013; Wu & Voeltz, 2021)*. The expression of the membrane-tethering modules SEC62_TMD_ (**Fig. 1D** and **Movie S3**), or TEX264_TMD_ (**Fig. 1E** and **Movie S4**) does not induce tubulation or constriction of the ER membrane.

### RHDs and TMDs determine the distribution of ER-phagy receptors in distinct ER subdomains

To determine the ER subdomain distribution of FAM134B_RHD_, SEC62_TMD_ and TEX264_TMD_, the three membrane-tethering modules were chemically cross-linked *in cellula* to adjacent endogenous polypeptides with 1 mM DSP. The immunocomplexes were isolated from cell lysates using GFP-trap nanobody-conjugated beads, and the interacting polypeptides were identified by Liquid Chromatography/Mass Spectrometry (LC/MS). FAM134B_RHD_ interacts with endogenous FAM134 family members, with proteins of the Reticulons, REEP and Atlastin families (green in **Figs. 1F, 1G**), hinting at a localization in high curvature ER subdomains. This is consistent with the reported subcellular distribution of full-length FAM134B, which localizes in regions of high ER curvature and controls their lysosomal clearance (Gonzalez *et al*., 2023; Khaminets *et al*., 2015).

In sharp contrast, SEC62_TMD_ (**Fig. 1F**) and TEX264_TMD_ (**Fig. 1G**) preferentially locate in rough ER subdomains characterized by the presence of sheet ER-forming proteins, molecular chaperones and enzymes of the protein disulfide isomerase and peptidyl:prolyl isomerase superfamilies, as well as proteins regulating ER translocation and *N*-glycosylation (red in **Figs. 1F, 1G**). This is consistent with the ER subdomain distribution of full-length SEC62 and TEX264, which ensure lysosomal turnover of portions of rough ER sheets containing folding chaperones and of the outer nuclear membrane (An *et al*, 2019; Chino *et al*, 2019; Fumagalli *et al*, 2016; Kucinska *et al*, 2023; Loi *et al*, 2018; Loi & Molinari, 2020; Loi *et al*, 2019). Thus, the membrane-tethering modules of FAM134B, SEC62 and TEX264 recapitulate the distinct intra-compartmental distribution reported for the full-length ER-phagy receptors (An *et al*., 2019; Chino *et al*., 2019; Fumagalli *et al*., 2016; Gonzalez *et al*., 2023; Khaminets *et al*., 2015; Kucinska *et al*., 2023; Loi *et al*., 2018; Loi & Molinari, 2020; Loi *et al*., 2019).

### Expression of reticulon-type and transmembrane-type ER-phagy receptors triggers ER-phagy

Ectopic expression of ER-phagy receptors of the FAM134 family recapitulates induction of ER-phagy as induced by nutrient-deprivation (Foronda *et al*., 2023; Gonzalez *et al*., 2023; Jiang *et al*., 2020; Khaminets *et al*., 2015; Reggio *et al*., 2021). However, there is some controversy in the available literature on the capacity of recombinant receptors lacking the RHDs that remodels the ER membrane to fragment the ER and recapitulate ER-phagy (Gubas & Dikic, 2022; Hoyer & Harper, 2023). To clarify this, we assessed the induction of ER-phagy in cells expressing the membrane-tethering modules of FAM134B, SEC62 and TEX264, or the same modules displaying their cytoplasmic IDRs engaging or not engaging LC3. All polypeptides were tagged at the C-terminus with HaloTag (HALO) (England *et al*, 2015), a bacterial alkane dehalogenase, whose active site has been modified to covalently bind cell-permeable fluorescent ligands (chloroalkane conjugated to tetramethyl-rhodamine (TMR) in our experiments).

Mock-transfected cells show a constitutive level of ER-phagy as quantified with the deep learning tool LysoQuant (Morone *et al*, 2020) in cells exposed to bafilomycin A1 (BafA1) to inhibit lysosomal hydrolases and preserve delivered material in the lumen of degradative organelles (Klionsky *et al*, 2008) (**Fig. 2A** and quantifications in **2E, 2I, 2M**, Mock). The expression of FAM134B_RHD_ does not trigger ER-phagy (**Figs. 2B, 2E**). As expected, recapitulation of ER-phagy requires that the RHDs of FAM134B displays the cytoplasmic IDR module that engages LC3 (FAM134B, **Figs. 2C, 2E**). The expression of FAM134B_LIR_ that carries a _453_DDFELL_458_ to _453_AAAAAA_458_ substitution that inactivates LC3 engagement (Khaminets *et al*., 2015) fails to enhance the delivery of ER portions within LAMP1-positive endolysosomes (**Figs. 2D, 2E**). Notably, the expression of SEC62 and TEX264, whose membrane-anchoring modules do not remodel the ER membrane (**Fig. 1**), potently activates lysosomal delivery of ER portions as well (**Figs. 2G, 2I** and **2K, 2M**), which is substantially inhibited by mutations of the LIRs that prevent LC3 engagement (SEC62_LIR_, **Figs. 2H-2I**, TEX264_LIR_, **Figs. 2L-2M**). Thus, the expression of ER-phagy receptors displaying an active IDR at the cytoplasmic face of the ER membrane triggers ER fragmentation and lysosomal delivery independent of the propensity of the ER-phagy receptor’ membrane-tethering module to remodel the ER membrane.

**Figure 2:**
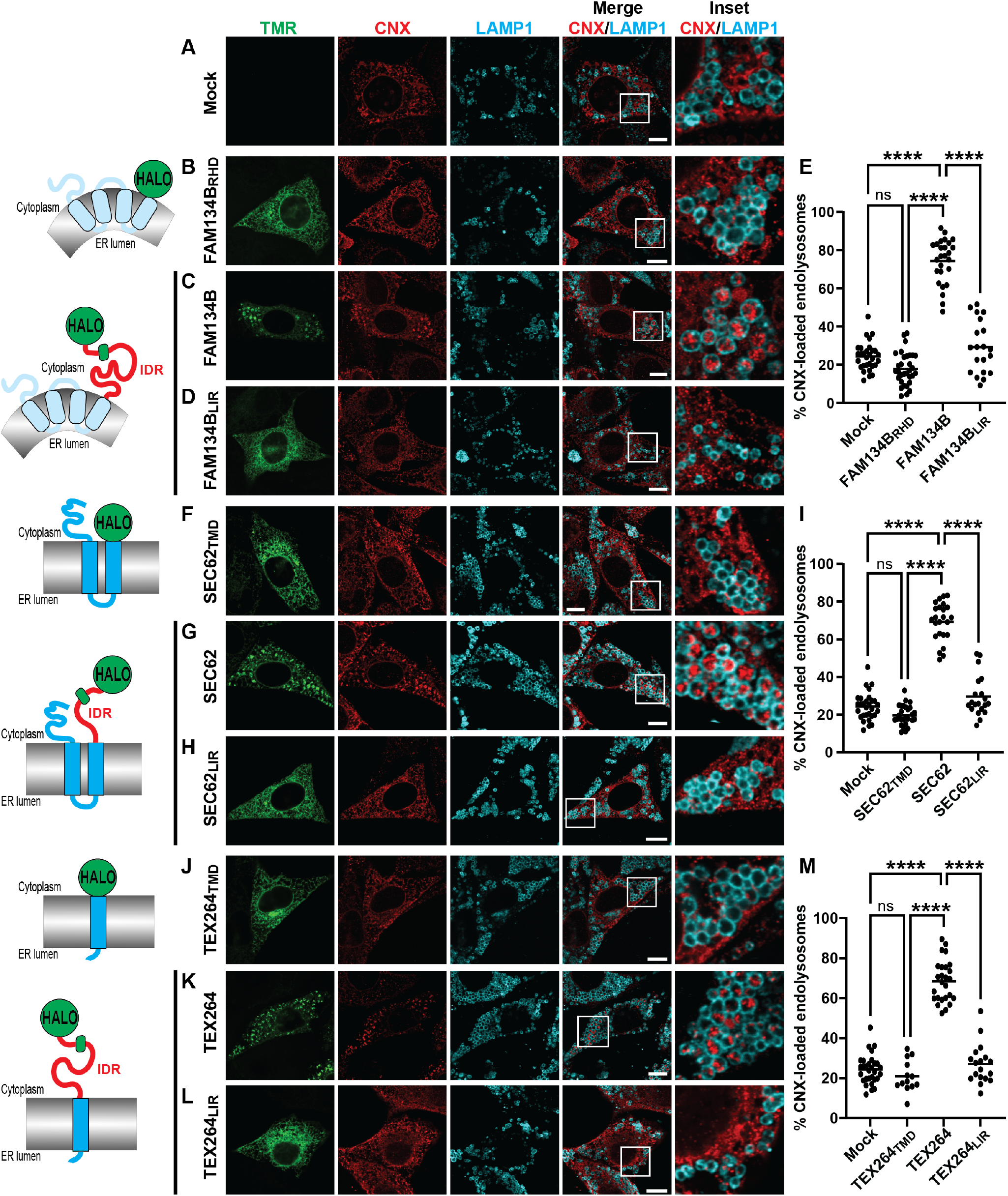
CLSM analyses of ER-phagy induction. **(A)** CLSM images of delivery of the ER marker CNX (red) to LAMP1-positive EL (cyan) in mock-transfected MEF treated with 50 nM BafA1 and 100 nM TMR. **(B)** Same as (A) in cells expressing FAM134B_RHD_-HALO (green, TMR). Scale bar: 10 μm; Inset: 4× magnification of the merge image. **(C)** Same as (B) in cells expressing FAM134B-HALO. **(D)** Same as (C) for FAM134B_LIR_-HALO. **(E)** LysoQuant quantification of the percentage of CNX-loaded LAMP1-positive EL in (A-D) (Mock [n=30 cells], FAM134B_RHD_-HALO [n=32 cells], FAM134B-HALO [n=26 cells], FAM134B_LIR_-HALO [n=20 cells]. N=3 biological replicates). **(F-H)** Same as (B-D) in cells expressing SEC62_TMD_-HALO, SEC62-HALO and SEC62_LIR_-HALO. **(I)** Same as (E) for SEC62_TMD_ -HALO [n=27 cells], SEC62-HALO [n=24 cells] and SEC62_LIR_-HALO [n=20 cells], N=3 biological replicates. Mock as in (E). **(J-M)** Same as (F-I) in cells expressing TEX264_TMD_-HALO [n=13 cells], TEX264-HALO [n=27 cells] and TEX264_LIR_-HALO [n=18 cells]. N=3 biological replicates. One-way ANOVA, F=190.1, 193.6 and 168 for (E), (I) and (M) respectively, ****P<0.0001, ns: not significant (P>0.05). Mean bar is shown.

### Expression of membrane-tethered IDRs elicits ER-phagy

Are membrane-bound IDRs engaging LC3 all what is needed to drive ER partitioning and capture by degradative organelles? To check this, we expressed in MEF the cytoplasmic IDRs of FAM134B, SEC62 or TEX264 (red in the schematics, **Fig. 3**) anchored at the ER membrane with a single spanning transmembrane domain of 17 residues (TM17) (Pedrazzini *et al*, 1996). The chimeras were tagged with a luminal monomeric sfGFP (**Fig. 3**). Quantitative analyses by CLSM show that the expression of a control sfGFP targeted to the ER membrane does not drive ER-phagy (**Fig. 3A**), while the expression of all ER-phagy receptor’s IDRs triggers ER fragmentation and capture of ER portions within LAMP1-positive endolysosomes (**Figs. 3B, 3D** for GT_17_-FAM134B_IDR_, **3E, 3G** for GT_17_-SEC62_IDR_, **3H, 3J** for GT_17_-TEX264_IDR_). The inactivation of the LC3-binding function abolishes delivery of ER portions within the LAMP1-positive endolysosomes (**Figs. 3C, 3D, 3F, 3G, 3I, 3J**).

**Figure 3:**
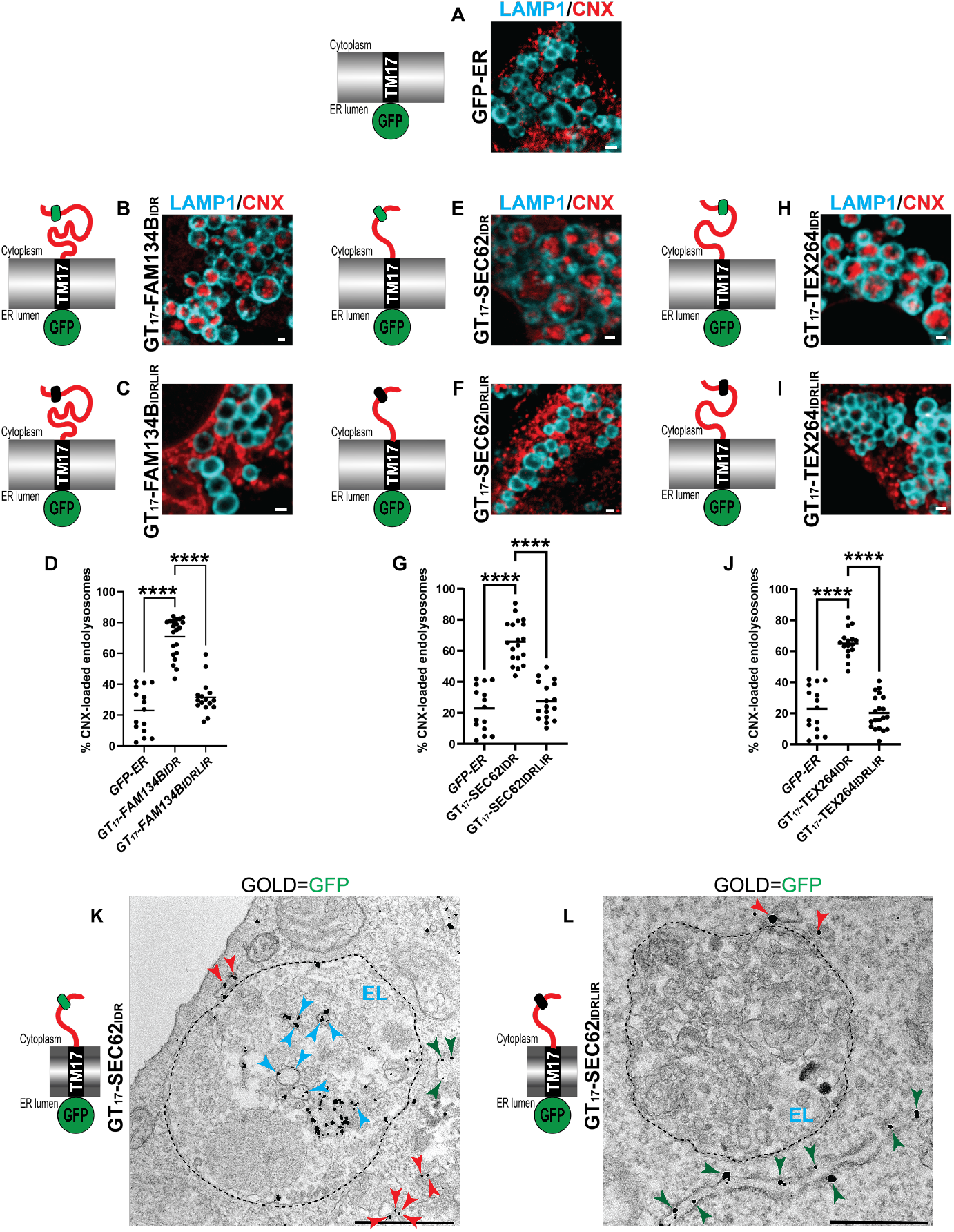
Membrane-associated IDRs trigger ER-phagy. Schematics of GT_17_-FAM134B_IDR_, GT_17_-SEC62_IDR_, and GT_17_-TEX264_IDR_ chimeras and corresponding LC3 binding-deficient (LIR) mutants are shown. **(A)** CLSM images of delivery of the ER marker CNX (red) to LAMP1-positive EL (LAMP1, cyan) upon the expression of GFP-ER control plasmid in MEF treated with 50 nm BafA1. **(B)** same as (A) for GT_17_-FAM134B_IDR_. **(C)** Same as (B) for GT_17_-FAM134B_IDRLIR_. Scale bars: 1 μm. **(D)** LysoQuant quantification of CNX-loaded LAMP1-positive EL in (A) [n=15 cells], (B) [n=21 cells] and (C) [n=16 cells], N=3 biological replicates. **(E-G)** Same as (B-D) for GT_17_-SEC62_IDR_ [n=19 cells] and GT_17_-SEC62_IDRLIR_ [n=_17_ cells], N=3 biological replicates. GFP-ER quantification is the same as in (D). **(H-J)** Same as (B-D) for GT_17_-TEX264_IDR_ [n=16 cells] and GT_17_-TEX264_IDRLIR_ [n=19 cells]. N=3 biological replicates. One-way ANOVA, ****P<0.0001, mean bar is shown. F=74.52 (D), 54.76 (G) and 78.4 (J). **(K)** IEM showing the localization of gold-labeled GT_17_-SEC62_IDR_ at the membrane of the ER (green arrowheads), of ER-derived vesicles (red arrowheads) and of ER-derived vesicles within EL (cyan arrowheads). Dashed line: EL limiting membrane. Scale bar: 500 nm. **(L)** Same as (K) for GT_17_-SEC62_IDRLIR_.

The competence of membrane-tethered IDR modules to fragment the ER and to activate ER capture by endolysosomes (**Figs. 3B-3J**) was also monitored with anti-GFP immunoelectron microscopy (IEM) in MEF treated with BafA1, where gold particles reveal the localization of GT_17_-SEC62_IDR_ at the ER membrane (**Fig. 3K**, green arrowheads), at the limiting membrane of cytoplasmic ER-derived vesicles (**Fig. 3K**, red arrowheads), and at the limiting membrane of ER-derived vesicles within endolysosomes (**Figs. 3L**, blue arrowheads and **3E**). GT_17_-SEC62_IDRLIR_, the mutant form that cannot engage LC3, also localizes in the ER (**Fig. 3L**, green arrowheads) and at the limiting membrane of cytoplasmic ER-derived vesicles (**Fig. 3L**, red arrowheads), but fails to deliver the ER portions within the endolysosomes (**Figs. 3L**, EL, and **3F**).

### Sub-compartmental distribution is lost when IDRs are tethered at the ER membrane with a generic TMD

Individual ER-phagy receptors localize and control lysosomal turnover of distinct ER subdomains (An *et al*., 2019; Chino *et al*., 2019; Fumagalli *et al*., 2016; Khaminets *et al*., 2015; Kucinska *et al*., 2023; Mochida *et al*, 2015; Reggiori & Molinari, 2022). The sub-compartmental distribution of the ER-phagy receptors is determined by the intrinsic properties of their membrane-anchoring modules with the RHDs distributing in ER subdomains enriched with proteins of the Reticulons, REEP and Atlastin families, whereas TMDs preferentially locate in rough ER subdomains characterized by the presence of sheet ER-forming proteins (**Figs. 1F, 1G**). This selectivity in subdomain distribution that determines the portion of the ER to be fragmented and delivered to the lysosomal compartment is lost upon replacement of the original membrane-tethering modules with a generic hydrophobic transmembrane sequence. In fact, the expression of FAM134B, SEC62 and TEX264 IDRs triggers the delivery to endolysosomes of both ER sheets (CLIMP63 in **Fig. 4**) and ER tubuli markers (REEP5 in **Fig. 5**).

**Figure 4:**
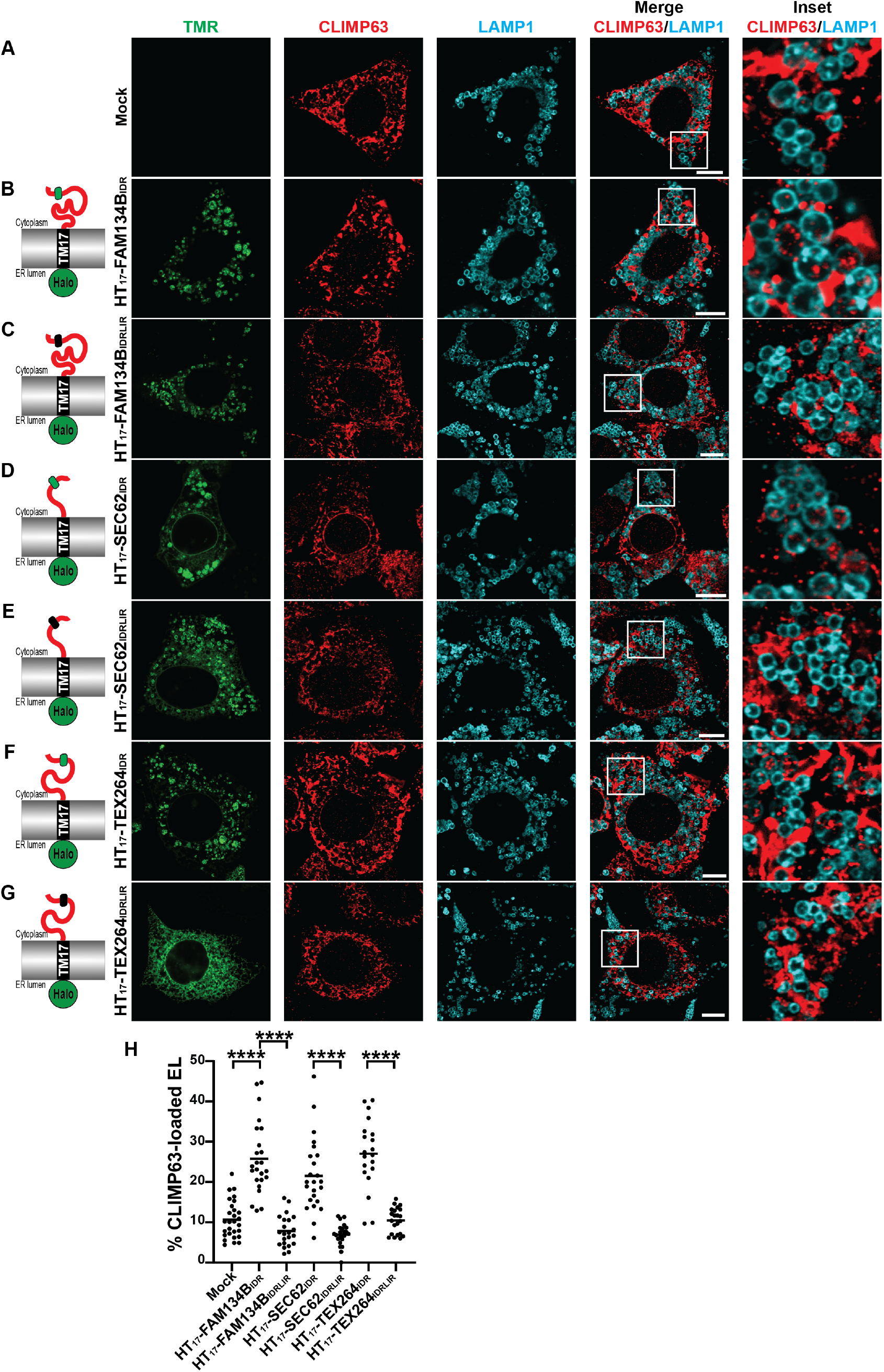
Expression of the IDR modules of FAM134B, SEC62 and TEX264 enhances endolysosomal delivery of CLIMP63. **(A)** CLSM images of delivery of the ER sheet marker CLIMP63 to LAMP1-positive EL in mock-transfected MEF treated with 50 nM BafA1 and 100 nM TMR. **(B)** Same as (A) in cells expressing HT_17_-FAM134B_IDR_ (TMR). Scale bar: 10 μm; Inset: 4× magnification of the merge image. **(C)** Same as (B) for the LC3-binding deficient HT_17_-FAM134B_IDRLIR_. **(D** and **E)** Same as (B and C) for the SEC62_IDR_ chimeras. **(F** and **G)** Same as (D and E) for the TEX264_IDR_ chimeras. **(H)** LysoQuant quantification of the percentage of CLIMP63-loaded LAMP1-positive EL in (A-G) (Mock [n=28 cells], HT_17_-FAM134B_IDR_ [n=25 cells], HT_17_-FAM134B_IDRLIR_ [n=24 cells], HT_17_-SEC62_IDR_ [n=26 cells], HT_17_-SEC62_IDRLIR_ [n=25 cells], HT_17_-TEX264_IDR_ [n=22 cells], HT_17_-TEX264_IDRLIR_ [n=22 cells], N=3 biological replicates). One-way ANOVA, ****P<0.0001, F=41.46. Mean bar is shown.

**Figure 5:**
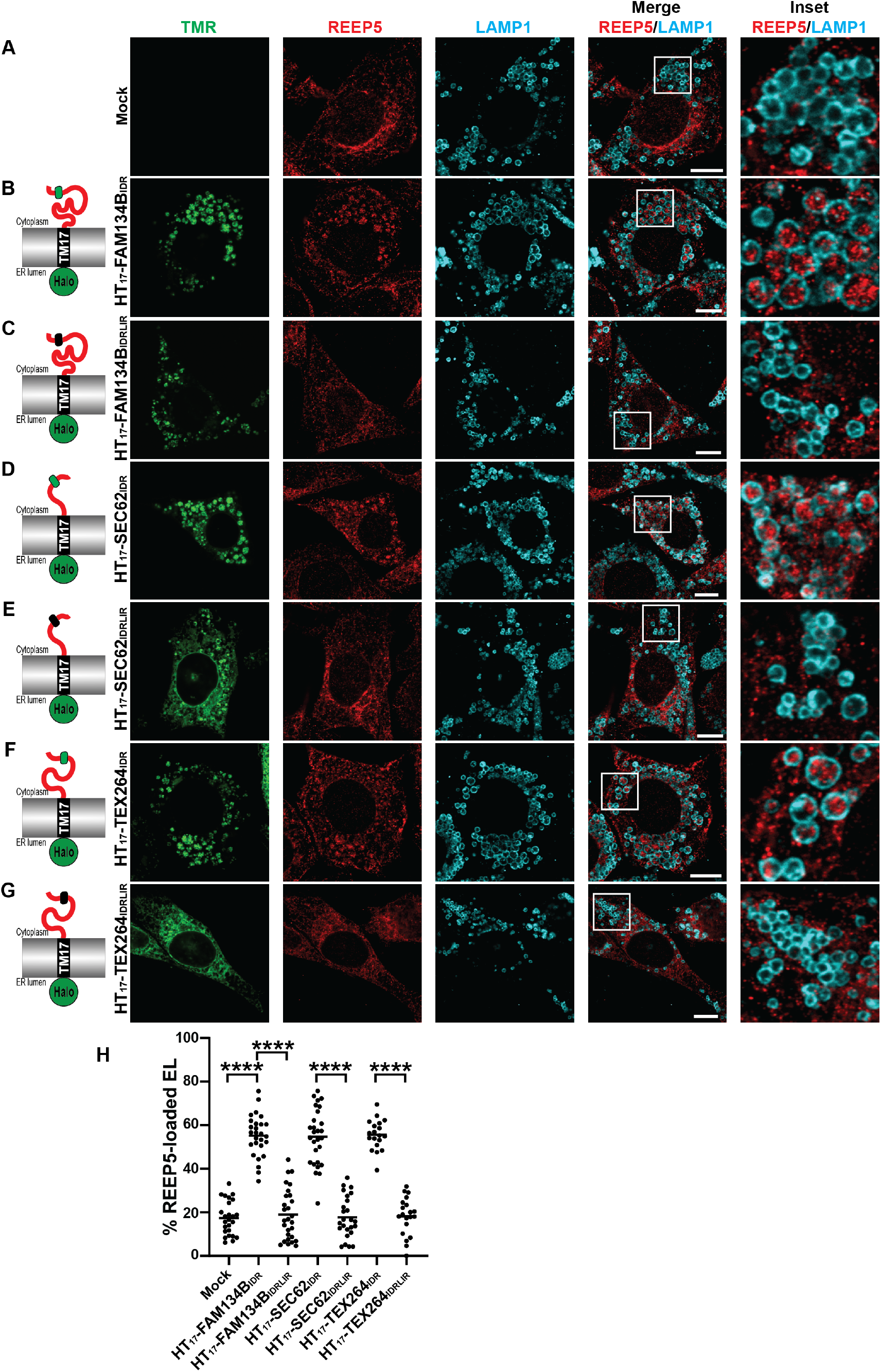
Expression of the IDR modules of FAM134B, SEC62 and TEX264 enhances endolysosomal delivery of REEP5. (**A**) CLSM images of delivery of the ER tubule marker REEP5 (red) to LAMP1-positive EL (cyan) in mock-transfected MEF treated with 50 nM BafA1 and 100 nM TMR. **(B)** Same as (A) in cells expressing HT_17_-FAM134B_IDR_ (green, TMR). Scale bar: 10 μm; Inset: 4× magnification of the merge image. **(C)** Same as (B) for the LC3-binding deficient HT_17_-FAM134BIDRLIR. **(D** and **E)** Same as (B and C) for the SEC62_IDR_ chimeras. **(F** and **G)** Same as (D and E) for the TEX264_IDR_ chimeras. **(H)** LysoQuant quantification of the percentage of REEP5-loaded LAMP1-positive EL in (A-G) (Mock [n=25 cells], HT_17_-FAM134B_IDR_ [n=27 cells], HT_17_-FAM134B_IDRLIR_ [n=28 cells], HT_17_-SEC62_IDR_ [n=27 cells], HT_17_-SEC62_IDRLIR_ [n=27 cells], HT_17_-TEX264_IDR_ [n=19 cells], HT_17_-TEX264_IDRLIR_ [n=20 cells], N=3 biological replicates). One-way ANOVA, ****P<0.0001, F=96.81. Mean bar is shown.

### LC3 lipidation is dispensable for IDRs-induced ER fragmentation

Delivery of ER portions within degradative organelles upon activation of ER-phagy by nutrient deprivation (An *et al*., 2019; Chino *et al*., 2019; Khaminets *et al*., 2015), recovery from ER stress (Fumagalli *et al*., 2016; Kucinska *et al*., 2023), or accumulation of misfolded proteins (Fregno *et al*, 2018) is abolished in MEF lacking ATG7, which are not lipidating LC3. Monitoring the ER ultrastructure upon expression of GT_17_-SEC62_IDR_ in MEF lacking ATG7 reveals the substantial ER remodeling (**Figs. 6C**), whereas the same construct lacking the cytoplasmic IDR fails to remodel the ER (**Fig. 6B**). Notably, the portions of ER generated by the IDR module cannot be delivered within the endolysosomal degradative compartments. Significantly, the same ER remodeling is observed upon expression of or GT_17_-SEC62_IDRLIR_ (**Figs 6D**) showing that the LIR, which is required to deliver ER portions within the LAMP1-positive compartment (**Fig. 3**), is dispensable for ER fragmentation.

**Figure 6:**
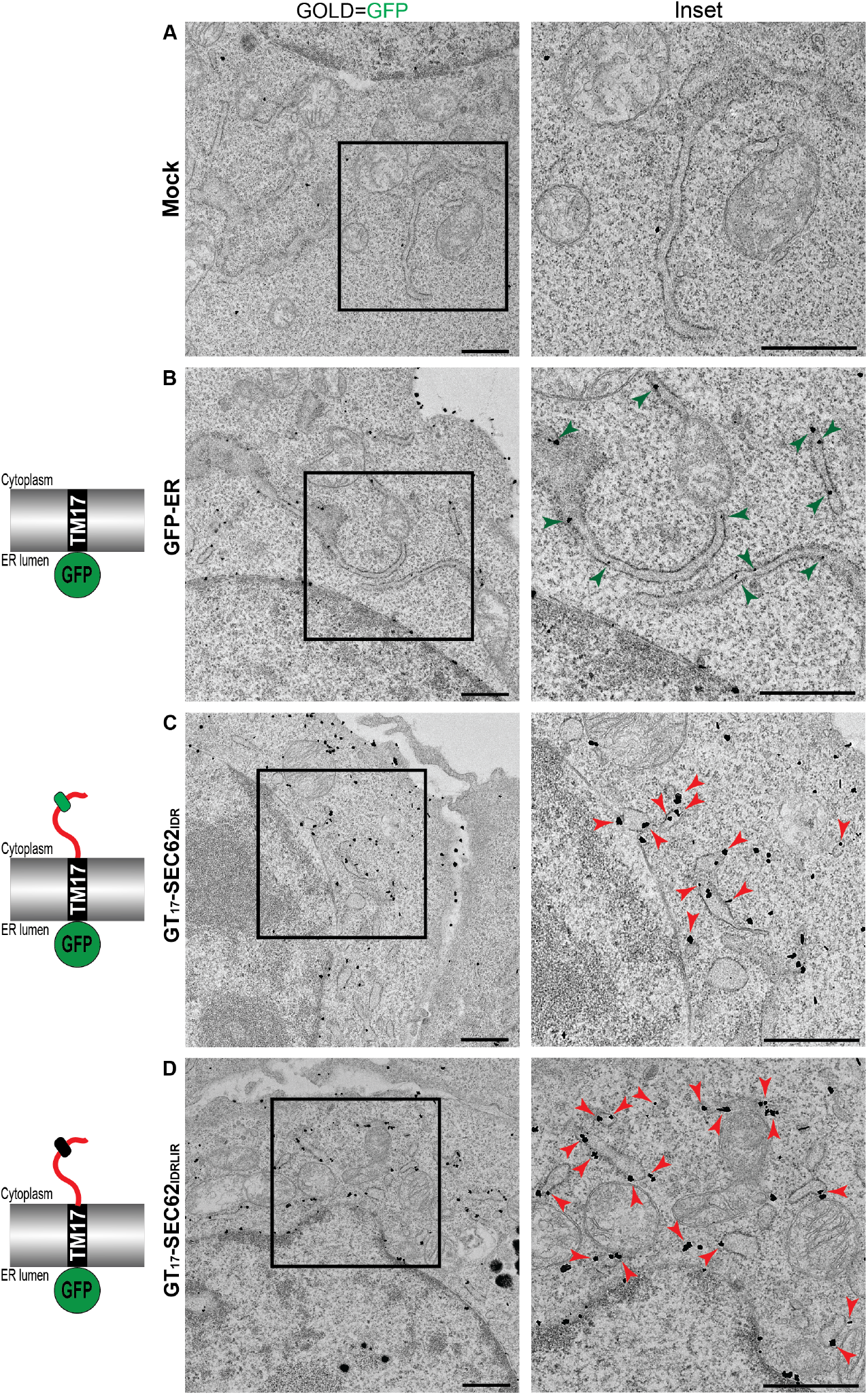
Expression of SEC62 IDR modules in ATG7-KO MEF triggers ER fragmentation. **(A)** IEM micrograph showing normal ER morphology in mock-transfected ATG7-KO MEF **(B)** Same as (A) in a cell expressing GFP-ER. Green arrows show gold-labelled ER sheets. **(C)** IEM micrograph showing fragmented ER in a cell expressing GT_17_-SEC62_IDR_. Red arrows indicate cytoplasmic ER fragments. **(D)** Same as (C) in a cell expressing GT_17_-SEC62_IDRLIR_. Scale bars: 1 μm, Inset magnification 2x.

## Discussion

Pleiotropic or ER-specific cues determine the lysosomal/vacuolar clearance of ER portions, including the adjacent nuclear envelope, to gain nutrients, to eliminate from cells toxic or aged macromolecules, to drive cell differentiation. These catabolic programs are driven by activation of ER-phagy receptors embedded within or recruited at the ER membrane to promote organelle fragmentation, a *conditio sine qua non* for execution of the ER-phagy programs that proceed upon engulfment of ER portions by the phagophore (*macro*-ER-phagy), upon direct engulfment of ER portions by degradative endolysosomes (*micro*-ER-phagy), or upon fusion of ER-derived vesicles with degradative endolysosomes (LC3-mediated delivery) (Reggiori & Molinari, 2022).

All ER-phagy receptors engage LC3 proteins via cytosolic exposed IDR modules, and our study proves that IDR modules of ER-phagy receptors are sufficient and required to promote ER fragmentation. The competence of the IDRs to engage LC3 ensures that the ER fragments are eventually delivered to the endolysosomal compartment for clearance. In line with our model, in a manuscript submitted to bioRxiv during preparation of our submission, molecular dynamics simulations reveal that the IDR module of FAM134B greatly amplifies the membrane remodelling activity of the RHD (Poveda-Cuevas *et al*, 2024) that was initially proposed to be sufficient to elicit ER-phagy (Bhaskara *et al*., 2019; Gonzalez *et al*., 2023). In an additional submission to bioRxiv, the group of Tom Rapoport reports on REEP1 that differs from other members of the reticulon homology domain containing ER -shaping proteins by the presence of a C-terminal disordered region that confers the peculiar capacity to generate ER-derived vesicles (Shibata *et al*, 2023). Thus, IDRs that, in the case of ER-phagy receptors have initially been proposed to bridge the distance between the ER membrane of the rough ER subdomains covered by ribosomes where the LC3-binding receptors are located, and the phagophore or the endolysosome membrane where LC3 is expected to be lipidated (Chino *et al*., 2019) play relevant roles in membrane remodelling. Not unexpectedly, long cytoplasmic IDRs are a feature conserved in ER-phagy receptors from yeast to mammals and by autophagy receptors exposed at the membrane of other organelles such as mitochondria and the Golgi compartment that need fission to enter the autophagic route and where there is no large steric hinderance to be expected such as that found in rough ER subdomains. Our data reveal a minimal distance of the LIR from the end of the membrane-tethering domain of about 24 residues to maintain the capacity to deliver ER portions within endolysosomes, which correlates with the competence to bind LC3 (**Fig. 7**). Again, consistent with a functional conservation of the IDR modules, a survey of the literature shows that for known membrane-bound organello-phagy receptors, the shortest distance of a LIR domain from the lipid bilayer are the 26 residues of the mitophagy receptor FUNDC1 (Liu *et al*, 2012). Considering the disordered nature of cytosolic LIR-containing peptide stretches, the presence of 24-26 amino acids would separate the LIR domain for about 9.5 nm from the lipid bilayer (Zhou, 2004), which is sufficient to accommodate LC3 molecules. All in all, the combination of a membrane-tethering module that specifies the sub-organellar portion to be removed from cells and of a functionally conserved IDR module that triggers organelle fragmentation and engages cytosolic autophagy gene products to deliver organelle portions to the degradative compartments builds an elegant and modulable effector to adapt organelle size and function to cellular needs.

**Figure 7:**
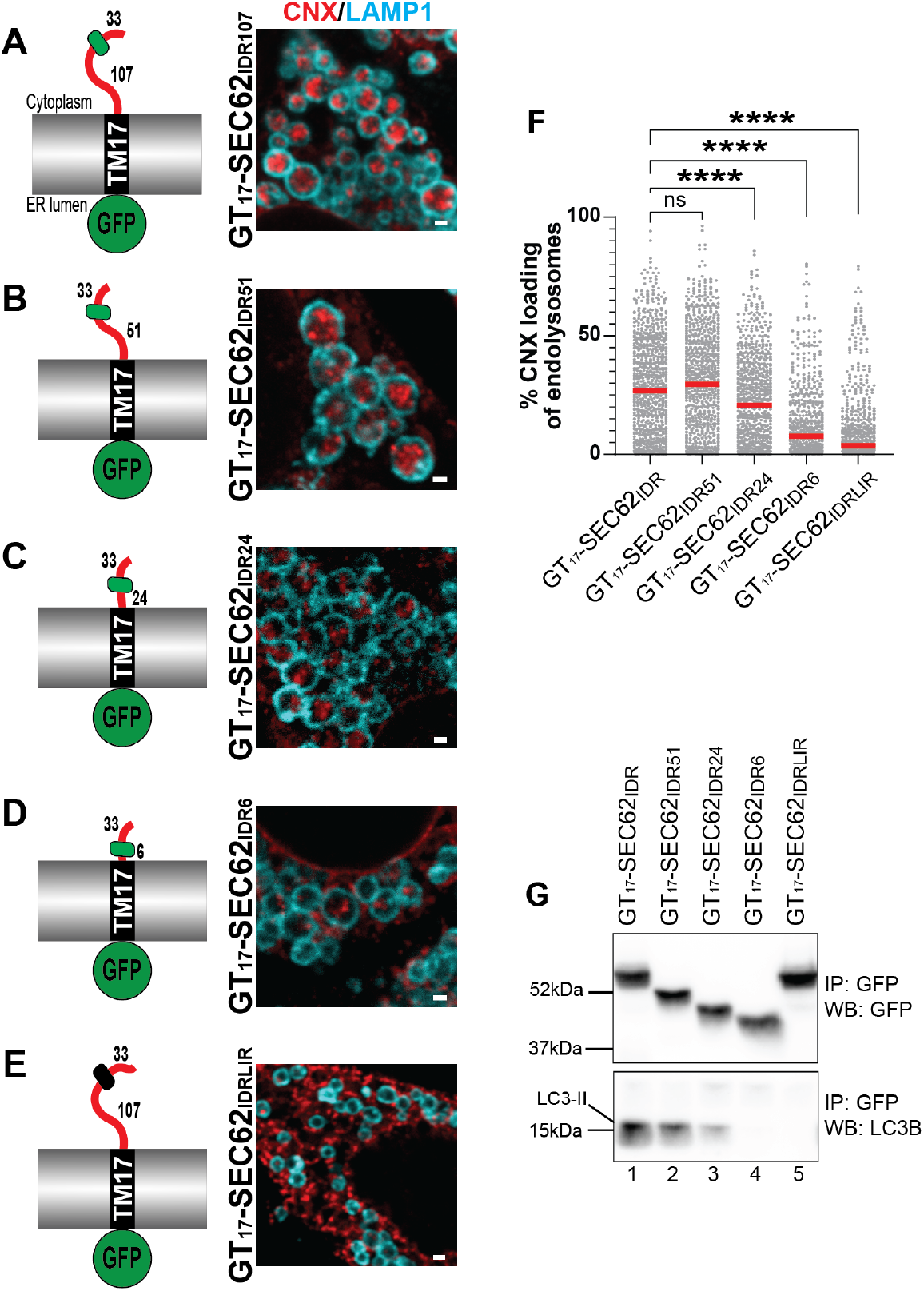
LC3 binding and ER-phagy by IDRs modules with progressively shorter distance between ER membrane and LIR. Schematics of GT_17_-SEC62_IDR_ chimeras of different IDR length and the LC3 binding-deficient GT_17_-SEC62_IDRLIR_. Numbers represent the length in the amino acids between the TM_17_ anchor and the LC3-interacting region, and between the LC3-interacting region and the C-terminus. **(A-E)** CLSM images of delivery of the ER marker Calnexin (CNX, red) to LAMP1-positive EL (LAMP1, cyan) upon the expression of GT_17_-SEC62_IDR_, _IDR51_, _IDR24_, _IDR6_ and GT_17_-SEC62_IDRLIR_ chimeras in MEF treated with 50 nm BafA1. Scale bars: 1 μm **(F)** LysoQuant quantification of the area of LAMP1-positive EL occupied by CNX in MEF from (A-D) (each dot represents a single EL, n = 811, 831, 739, 629, and 615 lysosomes analyzed in cells from (A-D) respectively, N=2 biological replicates). Kruskal-Wallis ANOVA test with Dunn’s multiple comparison test. Adjusted ****P < 0.0001, ns: not significant (P>0.05). Median bar is shown. **(G)** Association of endogenous LC3B-II with GT_17_-SEC62_IDR_ chimeras in MEF treated with 50 nM BafA1. Immunocomplexes were revealed with anti-GFP (upper panel) or anti-LC3B antibodies (lower panel).

Notably, all membrane-bound and soluble ER-phagy receptors (in yeast, plant and mammalian cells) and all receptors regulating mito-phagy and Golgi-phagy display cytoplasmic IDR modules (Delorme-Axford *et al*, 2019; Hickey *et al*, 2023; Johansen & Lamark, 2020; Popelka & Uversky, 2022; Reggiori & Molinari, 2022). Organellophagy receptors’ IDRs lack a defined 3D structure and amino acid sequence homology (Lemke *et al*, 2024; van der Lee *et al*, 2014) but are all characterized by net negative charges (e.g., -19.84 for SEC62_IDR_, -24.16 for FAM134B_IDR_, -16.31 for TEX264_IDR_, -3.93 for mitochondrial FUNDC_IDR_, -11.96 for Golgi YIPF3_IDR_, -11.11 for Golgi YIPF4_IDR_, as calculated with https://www.biosynth.com/peptide-calculator). A survey of the literature reveals that the function of IDR includes the processing of regulatory cues, molecular communication (Holehouse & Kragelund, 2023) and, notably, sensing and driving membrane curvature and fission (Busch *et al*, 2015; Shibata *et al*., 2023; Snead *et al*, 2017; Zeno *et al*, 2018; Zeno *et al*, 2019). We anticipate that activation of the exposure of IDR modules at the limiting membrane of organelles could facilitate fragmentation and, upon LC3 association, lysosomal clearance.

## Supporting information

SMovie 1 Mock-transfected for Fig. 1B

SMovie 2 FAM134B-RHD for Fig. 1C

SMovie 3 SEC62-TMD movie for Fig. 1D

SMovie 4 TEX264-TMD movie for Fig. 1E

## Acknowledgments

We thank the members of Molinari’s laboratory and Y. Grishchuk for discussions, and critical reading of the manuscript.

## Author contributions

Conceptualization: MM, MR, CG

Methodology: MR, CG, AR, MM

Investigation: MR, CG

Visualization: MR, CG, AR, MM

Funding acquisition: MM

Project administration: MM

Supervision: MM

Writing – original draft: MM

Writing – review & editing: MM, MR, CG

## Data and materials availability

All data, code, and materials used in the analysis are available, without restriction upon reasonable request to reproduce or extend the analyses.

## Methods

### Cell Culture, Transient Transfection, and Inhibitors

Mouse embryonic fibroblast (MEF) and Human Embryonic Kidney 293 (HEK293) cells were cultured in DMEM supplemented with 10% fetal calf serum (FCS) at 37°C and 5% CO_2_. Transient transfections were performed using JetPrime transfection reagent (PolyPlus) following the manufacturer’s protocol. Bafilomycin A1 (BafA1, Calbiochem) was used at 50 nM for 15 hours if not otherwise specified. WT and ATG7-KO MEF are a kind gift from M. Komatsu.

### Expression Plasmids and Antibodies

ER-targeted IDR constructs were subcloned in a pcDNA3.1(+) backbone with an N-terminal GFP tag. IDR portion was separated from TM17 transmembrane anchor by a Tyr-Arg-Gly-Ser (YRGS) linker. The ER targeting was achieved by incorporating an N-terminal prolactin signal sequence. Constructs containing transmembrane domain modules were fused with a C-terminal GFP or HALO moiety. Antibodies used in this study are summarized in **Table S1**.

### Cell lysis and Western Blot

After the respective treatments, MEF were washed with ice-cold phosphate-buffered saline (PBS) containing 20 mM N-ethylmaleimide (NEM). Cells were lysed with either 2% CHAPS (in HEPES-buffered saline [HBS], pH 7.4) or RIPA buffer (1% Triton X-100, 0.1% SDS, 0.5% sodium deoxycholate in HBS, pH 7.4), supplemented with 20 mM NEM and protease inhibitors (1 mM phenylmethylsulfonyl fluoride (PMSF), 16.5 mM Chymostatin, 23.4 mM Leupeptin, 16.6 mM Antipain, 14.6 mM Pepstatin) for 20 min on ice. The postnuclear supernatant (PNS) was extracted by centrifugation at 4°C, 10,600 × *g* for 10 minutes. The PNS was denatured with the addition of 100 mM dithiothreitol (DTT) for 5 min at 95°C and subjected to SDS-PAGE. Proteins were then transferred to polyvinylidene difluoride (PVDF) membranes using the Trans-Blot Turbo Transfer System (Bio-Rad). The membranes were blocked with 8% (weight/vol) nonfat dry milk (Bio-Rad) in Tris-buffered saline with 1% Tween 20 (Sigma Aldrich) (TBS-T), and then incubated overnight with primary antibodies diluted in TBS-T. Following this, horseradish peroxidase (HRP)-conjugated Protein A diluted in TBS-T was applied for 45 min. Protein bands were visualized using the WesternBright Quantum detection system (Advansta), and the signals were captured using the FusionFX chemiluminescence imaging system (VILBER). The western blot bands were quantified using the FIJI software(Schindelin *et al*, 2012).

### Protein Cross-Linking with dithiobis-succinimidyl propionate (DSP)

Following treatment, cells were PBS-washed and treated with 1 mM DSP (Thermo Fischer Scientific) (from a 100x dimethylsulfoxide (DMSO) solution) in PBS. DSP incubation occurred for 30 min at room temperature (RT). The reaction was halted by adding 1 M Tris, pH 7.8 (final concentration 20 mM), and incubating for an additional 15 min at RT. Cells were then PBS-washed, treated with 20 mM NEM, and lysed with RIPA buffer (1% Triton X-100, 0.1% SDS, 0.5% sodium deoxycholate in HBS, pH 7.4) for 20 min on ice. After centrifugation at 10,600 g for 10 min, the post-nuclear supernatants (PNS) were collected and utilized for immunoprecipitation of IDR chimera-LC3 complexes.

### Affinity Purification-Liquid Chromatography/Mass Spectrometry (AP-LC/MS)

HEK293 cells were transfected with GFP-tagged TMD/RHD chimeras. Fifteen hours post-transfections cells were treated with 100 nM BafA1 for 6 hours, crosslinked with DSP as described above, and the supernatant was collected. Protein complexes were isolated with 100 µl GFP-TRAP (Chromotek) following manufacturer’s protocol, digested, desalted and injected into Orbitrap Exploris (Thermo Scientific). Intensity values were obtained following MaxQuant (Cox & Mann, 2008) analysis. Only the proteins with at least two detected peptides were considered for analyses. The results were normalized by median abundance, the missing values were imputed from a normal distribution of 1.8 standard deviation down shift and with a width of 0.3 of each sample, and unpaired two-tailed tests were used to calculate the p-value. Data analyses and graph plotting were done with the Perseus platform (version 2.0.5) (Tyanova *et al*, 2016) and MATLAB (The MathWorks Inc., R2023b Update 5 23.2.0.2459199 for Windows).

### Confocal Laser Scanning Microscopy (CLSM)

MEFs were seeded on alcian blue-treated (Sigma) glass coverslips (VWR) and transiently transfected using JetPrime reagent according to the manufacturer’s protocol. For Lysoquant analyses, 8 h after transfection, cells were treated with 50 nM BafA1 for 15 h. In addition, cells transfected with HALO-fusion protein were supplemented with 100 nM TMR HALO ligand (Promega). Cells were fixed at RT for 20 min in 3.7% formaldehyde (FA) diluted in PBS 24 h after transfection. Membrane permeabilization was performed by incubating the coverslips for 20 min in permeabilization solution (PS, 10% goat serum, 10 mM HEPES, 15 mM glycine, 0.05% saponin). Following permeabilization, cells were incubated with the primary antibodies diluted 1:100 in PS (unless otherwise specified in table S1) for 120 min, washed three times in PS, and then incubated with Alexa Fluor-conjugated secondary antibodies diluted 1:300 in PS for 45 min. Cells were rinsed three times with PS and water and mounted with a drop of Mount Liquid anti-fade (Abberior) on glass microscope slides (Epredia). Confocal images were acquired using a Leica TCS SP5, Leica STELLARIS 5 and Leica STELLARIS 8 microscopes equipped with Leica HCX PL APO lambda blue 63.0 × 1.40 oil objective with pinhole 1 AU. Leica LAS X software was used for image acquisition, with excitation provided by 489-, 499-, 552-, 561-, 587-, 641-, and 653-nm laser beans, and fluorescence emission collected within the ranges of 494-557 nm (GFP), 504-587 nm (AlexaFluor488), 557-663 nm (TMR), 592-644 nm (AlexaFluor568) and 658-750 nm (AlexaFluor646), respectively. Image analysis and quantification were performed with LysoQuant and FIJI software (Morone *et al*., 2020; Schindelin *et al*., 2012). Image post-processing was performed with Adobe Photoshop.

### Room Temperature-Transmission Electron Microscopy (RT-TEM) and Electron Tomography (RT-ET)

MEFs were plated on gridded glass bottom dishes (MatTek Corporation) and transiently transfected using JetPrime reagent following the manufacturer’s protocol. Twenty-four hours after transfection, cells were pre-fixed with a 4% FA EM-grade solution supplemented with 0.1% glutaraldehyde (GA). The coordinates of the cells on the finder grid were determined using wide-field microscopy on the Leica STELLARIS 8 microscope with Leica PL APO 10x0.40 air objective with pinhole 2 AU, and the transfected and non-transfected cells were identified based on GFP fluorescence (493-556 nm range). Cells were then fixed with a solution containing 2.5% GA in 0.1 M sodium cacodylate buffer (pH 7.4). After several washes in sodium cacodylate buffer, cells were post-fixed in 1% osmium tetroxide (OsO4), 1.5% potassium ferricyanide (K4[Fe(CN)6]) in 0.1 M sodium cacodylate buffer for 1 h on ice, washed with distilled water and stained *en bloc* with 0.5% uranyl acetate in distilled water overnight at 4°C in the dark. Samples were then rinsed in distilled water, dehydrated with increasing concentrations of ethanol, and embedded in Epoxy resin (Sigma Aldrich). After curing for 48 h at 60°C, ultrathin sections (70-90 nm thickness) were collected using an ultramicrotome (UC7, Leica microsystem), stained with uranyl acetate and Sato’s lead solutions, and observed under a Transmission Electron Microscope Talos L120C (FEI, Thermo Fisher Scientific) operating at 120 kV. Images of transfected and non-transfected cells were acquired with a Ceta CCD camera (FEI, Thermo Fisher Scientific). For 3D serial section electron tomography, serial sections (130-150nm thickness) were collected on formvar-carbon coated slot grids, tilted images series (+60°) were acquired with a Talos L120C TEM using Tomography 5 software (FEI, Thermo Fisher Scientific). Tilted images were aligned, and tomograms reconstructed using IMOD software (v. 4.11.24) (Organelle segmentation was performed in Microscopy Image Browser (MIB, v. 2.84 (Belevich *et al*, 2016)) and visualized in IMOD (Kremer *et al*, 1996)).

### Immunogold Electron Microscopy (IEM)

MEF were plated on Nunc Thermanox plastic coverslips (Thermo Fisher Scientific) and transiently transfected with GFP-tagged chimeras using the JetPrime reagent following the manufacturer’s protocol. Eight hours post-transfection, cells were either treated with 50 nM BafA1 for 15 h and fixed for 2 h in periodate-lysine-paraformaldehyde (PLP, (McLean & Nakane, 1974) FUJIFILM Wako) at RT or directly fixed 25 h after transfection. After fixative washout with PBS 1X, cells were incubated with 50 mM glycine and blocked for 30 min in blocking buffer (0.2% bovine serum albumin, 5% goat serum, 50 mM NH4Cl, 0.1% saponin, 20 mM PO4 buffer, 150 mM NaCl) at RT. Staining with primary rabbit anti-GFP antibody (Abcam) and gold-labeled secondary antibodies (Nanoprobes) was performed in a blocking buffer at RT. Cells were refixed for 30 min in 1% GA, and the nanogold particles were enlarged with gold enhancement solution (Nanoprobes) according to the manufacturer’s instructions. Cells were post-fixed with OsO4 and processed as described for EM. Images were acquired with a Ceta CCD camera (FEI, Thermo Fisher Scientific) using Velox 3.6.0 (FEI, Thermo Fisher Scientific) on Talos L120C TEM (FEI, Thermo Fisher Scientific) operating at 120 kV.

### Statistical analyses

The statistical comparisons and graphical plots were performed using GraphPad Prism 10 (GraphPad Software Inc., version 10.1.2 for Windows) and MATLAB (The MathWorks Inc., R2023b Update 5 23.2.0.2459199 for Windows). Tests employed for statistical analysis are reported in figure legends. An adjusted *P* value < 0.05 (for one-way ANOVA with multiple comparison tests) was considered statistically significant. Anova test results were corrected for multiple comparisons, and F values are reported in figure legends. All experimental replicates represent independent biological replicates. Figure legends specify the number of cells or lysosomes analyzed (n) and the number of independent biological replicates of the experiment (N).

**Table S1.** Details of antibodies and fluorescent ligands used

| Antibody | Manufacturer | Catalog number | Dilution or Concentration |
| --- | --- | --- | --- |
| Rat anti-LAMP1 | DSHB | 1D4B | 1:50 (CLSM) |
| Rabbit anti-LC3 | Sigma | L7543 | 1:1000 (IB) |
| Rabbit anti-CNX | Kind gift from A. Helenius | Not applicable | 1:100 (CLSM) |
| Rabbit anti-CLIMP63 | Proteintech | 16686-1-AP | 1:100 (CLSM) |
| Rabbit anti-REEP5 | Proteintech | 14643-1-AP | 1:100 (CLSM) |
| Rabbit anti-GFP | Abcam | ab290 | 1:50 (IEM) 1:1500 (IB) |
| GFP-TRAP | Chromotek | gta | Not applicable |
| Protein A HRP-conjugated | Invitrogen | 101023 | 1:20000 (IB) |
| Goat anti-rabbit AlexaFluor488-conjugated | Thermo Fisher Scientific | A-21206 | 1:300 (CLSM) |
| Goat anti-rabbit AlexaFluor568-conjugated | Thermo Fisher Scientific | A-11036 | 1:300 (CLSM) |
| Goat anti-rat AlexaFluor647-conjugated | Thermo Fisher Scientific | A-21247 | 1:300 (CLSM) |
| Goat anti-rabbit gold-labelled | Nanoprobes | 2004 | 1:100 (IEM) |
| HaloTag TMR Ligand | Promega | G8252 | 100 nM (CLSM) |

